# How can we study the evolution of animal minds?

**DOI:** 10.1101/024422

**Authors:** Maxime Cauchoix, Alexis Chaine

## Abstract

During the last 50 years, comparative cognition and neurosciences have improved our understanding of animal minds while evolutionary ecology has revealed how selection acts on traits through evolutionary time. We describe how this evolutionary approach can be used to understand the evolution of animal cognition. We recount how *comparative* and *fitness methods* have been used to understand the evolution of cognition and outline how these methods could be extended to gain new insights into cognitive evolution. The fitness approach, in particular, offers unprecedented opportunities to study the evolutionary mechanisms responsible for variation in cognition within species and could allow us to investigate both proximate (ie: neural and developmental) and ultimate (ie: ecological and evolutionary) underpinnings of animal cognition together. Our goal in this review is to build a bridge between cognitive neuroscientist and evolutionary biologists, illustrate how their research could be complementary, and encourage evolutionary ecologists to include explicit attention to cognitive processes in their studies of behaviour. We believe that in doing so, we can break new ground in our understanding of the evolution of cognition as well as gain a much better understanding of animal behaviour.

## Introduction

Niko Tinbergen (Tinbergen, 1963) proposed that biologists should try to understand animal behaviours in the light of two different and complimentary perspectives: the *proximate* and *ultimate* (see Bateson and Laland, 2013; Laland et al., 2011 for recent updates). While both approaches have been employed in the study of animal cognition, most studies have done so independently with little integration across fields. After some promising, integrative studies in the 1980s and 1990s (see Kamil, 1998 for a review), the last decades have seen the establishment of entirely independent lines of research with only a few notable exceptions. We now have a deeper understanding of how animal minds work, but we know very little about the evolution of or ecological pressures that shape cognition. Consequently, we know very little about what role cognition, a collection of highly plastic and flexible traits, plays in adaptation and biological evolution. We believe the time is ripe for evolutionary ecology studies to explicitly integrate cognition to generate a much stronger understanding of how the mind evolves.

Proximate studies focus on the mechanisms underlying given behaviours and the developmental biology of key structures. What stimuli trigger behaviours? How do neurons in the brain encode stimuli and transform them into behaviour? What is the ontogeny of behaviour? In other words, the proximate approach tries to understand how animal minds work. The current view for cognitive neuroscientists is that the animal mind emerges from brain activity as the neural machinery encodes, manipulates, stores and recalls information, which is together called ‘*cognition*’. Cognition emerges when the brain transforms information into mental constructs or representations (Barsalou, 2014). For *cognitive scientists*, cognition is a synonym of ‘mind’, which, operationally, is divided in various *cognitive functions*, each function being implied in a specific step of information processing (see also Figure 1). Perception (i.e. vision, olfaction, audition, gustation and somesthesia) all contribute to the process by which mental representations are built from environmental stimulation. Learning is the ability to associate previously unrelated mental representations. Memory is the ability to store mental representations either for a small amount of time (short term memory), a large amount of time (long term memory) or in relation to a particular on-going task (working memory). Attention is the mechanism allowing an individual to focus on only some mental representations among many. Decision-making is the process enabling an individual to compare mental representations and choose the most appropriate given the environmental context. Finally, executive functions (reasoning, problem solving, flexibility, categorization etc…) enable an individual to perform operations and manipulations of mental representations. Cognition is also sometimes divided according to the nature of the representation; one can for instance talk about spatial or social cognition.

**Figure 1:**
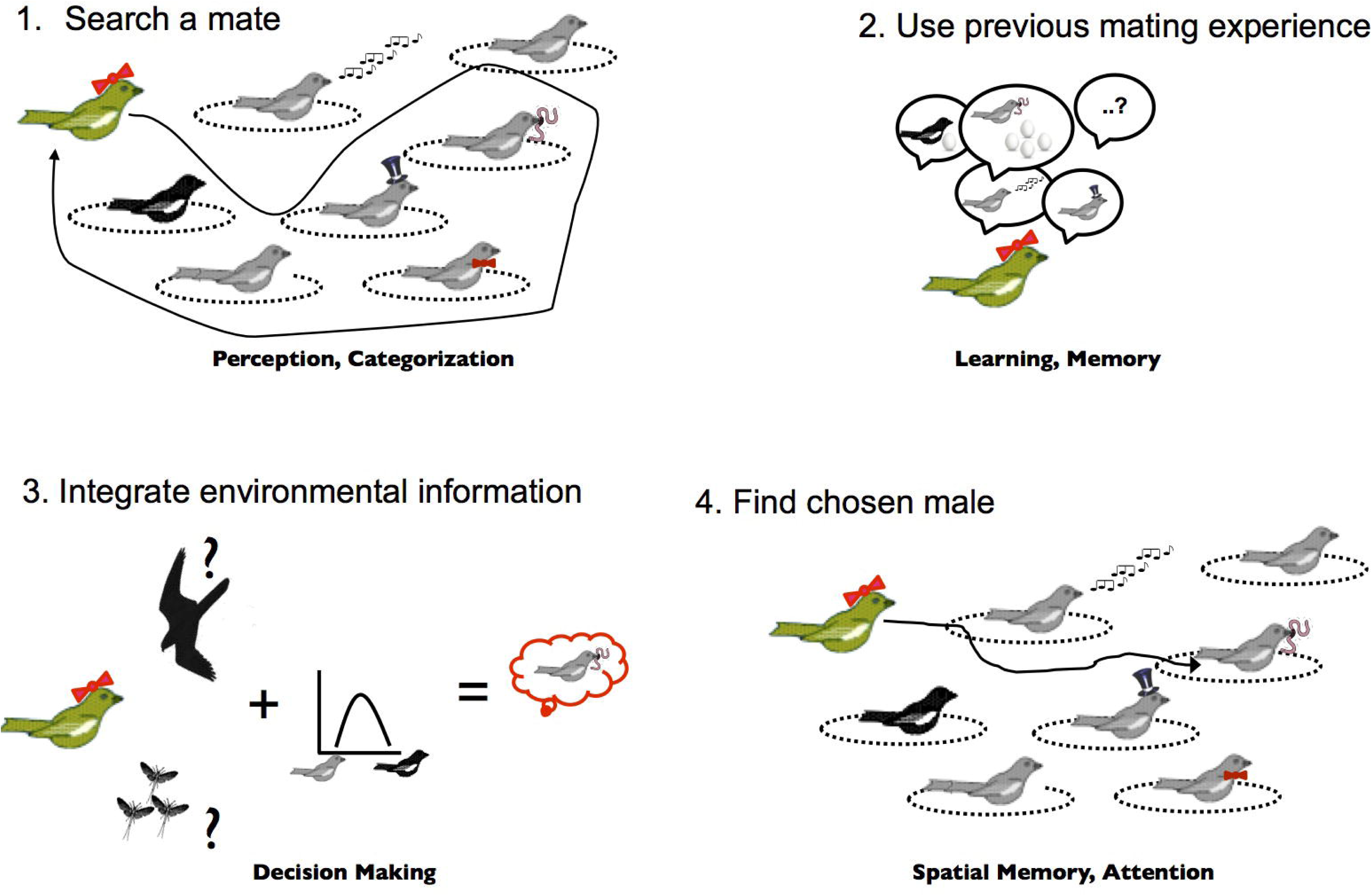
Mate choice and cognitive capacities that could hypothetically play a role. In bi-parental breeding songbirds, choosing an appropriate mate according to available male stock, previous breeding experience and actual environmental conditions is a behaviour that will have drastic fitness consequences for any female and that is likely to rely on the interplay between various cognitive functions. Recognition of ornaments linked to different male qualities (e.g. good genes, parental care, nest defense, etc.) uses perception (visual and auditory) to detect male signals and categorization to group and identify male quality according to their ornaments (1). The use of previous breeding experience relies on past learning linking male ornaments and reproductive success from previous experiences (2). Mate choice itself, integrates all information available to the female including current ecology, mate options, and past experience supposedly through decision-making mechanisms (3). Finding the chosen mate, once the decision has been taken, probably relies on spatial memory to relocate the territory defended by the chosen male and endogenous attention to detect the chosen male from among the background of other males and environmental features (4).

The association between studies in psychology and neurosciences along with the advent of powerful new neuroimaging technics (e.g. In vivo electrophysiology, Magnetic Resonance Imaging (MRI), Positron emission Tomography (PET), optogenetic etc.) has lead us to better understand how behaviours and decisions are linked to neural structures and neural activity in several animal species including humans. Despite this in depth understanding, much less progress has been made in understanding the evolutionary processes that have lead to the patterns of cognition that we see.

Ultimate approaches focus on the evolutionary history of behaviours or traits and the selective pressures that favour the evolution of those traits. Those using this approach have focused on behaviours with only a few rare studies examining cognition *per se* (e.g. Bond and Kamil, 2002, 2006; Lyon, 2003; Théry and Casas, 2002). *Evolutionary biologists* and *behavioural ecologists* have been primarily interested in the ecology and evolution of behaviour without examining the cognitive mechanisms underlying these behaviours. What ecological or social contexts are responsible for the evolution of a specific behaviour? What role does evolutionary history (inheritance from a common ancestor) play in the evolution of that trait? What are the costs and benefits of behaviours and what do they imply for selection on the animal’s life history strategy? To answer these questions behavioural ecologists have adopted the Neo-Darwinian theoretical framework and developed tools and models to understand the extreme variability of behaviours within and among species. However, this approach focuses on the aggregate outcome of cognition and action (i.e. the behaviour) and has usually considered the animal mind as a black box (Giraldeau, 2004). Indeed, much of behavioral or evolutionary ecology theory is based on strategic decision-making. While in some cases these strategic decisions reflect physiological trade-offs, many more cases reflect decisions made probably on the basis of processing external information gathered by an individual. Attention in such studies is placed on the quality of information and the outcome of a decision, but there is little understanding of how information is processed and how cognitive abilities enhance or constrain decisions based on the available information (Rowe, 1999, 2013). For example, social behavior, individual recognition, mate choice, parental care, dispersal, foraging, and predator avoidance nearly always rely on gathering external information. How well an individual gathers that information, how well it remembers that information, and how it integrates different sources of information all depend on cognitive capacities. To illustrate this notion (Figure 1) we can imagine a female who must choose the best mate among males that each display a number of ornaments linked to various qualities (e.g. good genes, parental care, nest defense, etc…). How does a female integrate the information provided in each of the male’s sexual signals with information about the external ecological environment (e.g. are there many nest predators)? As the female comparison shops for the best male, how many of the males can she remember? If she chooses to return to the second male she saw, will she remember where he is and will she recognize him? This example illustrates just some of the cognitive processes related to one behavior that would have fundamental consequences for sexual selection theory. Many other behaviors and life history strategies will similarly depend on cognitive capacities and actual measurement of cognitive abilities has the potential to fundamentally alter our views of behavior.

Understanding the evolutionary and ecological significance of cognition has been a major challenge in biology as highlighted in several recent books (Dukas and Ratcliffe, 2009; Heyes and Huber, 2000; Shettleworth, 2010) and review articles (Boogert et al., 2011; Dukas, 2004, 2008; Healy and Braithwaite, 2000; Kamil, 1998; MacLean et al., 2012; Pravosudov and Smulders, 2010; Real, 1993; Thornton et al., 2012) and has led to a new field of research called *cognitive ecology*. We argue that two factors will help to significantly advance our understanding of animal cognition: 1) proximate and ultimate studies should develop lines of research that allow direct integration of the two fields and 2) that evolutionary studies begin to apply their research methods to cognition *per se* along with the behaviours that result from cognitive processes. In doing so, we will gain a better understanding of how cognitive systems evolve and how cognitive structures and function relate to the problems they evolved to solve.

In this review, we focus more on the contribution that evolutionary biology can offer cognitive research since much less work has been done in this domain. Despite this bias towards what evolutionary biologists could contribute (i.e. what we know less about), we also highlight new contributions that cognitive neuroscientists could make to better integrate proximate and ultimate understandings of cognition. In the first section, we review past work testing popular hypotheses for cognitive evolution using comparative methods and highlight future directions to exploit using these methods. We then illustrate how measuring selection on cognition within a species provides a great opportunity to better understand the evolution of cognition and create direct links with proximate studies of cognition (e.g. neurosciences, cognitive-psychology). We finish by presenting two lines of research as case studies—food hoarding and brood parasitism—that, in our view, have best integrated ecological challenges, natural behaviour and underlying cognitive adaptation and which could serve as examples for future cognitive ecology research.

### Phylogenetic comparative studies of cognitive evolution

Current tests of factors that influence the evolution of the brain have largely relied on comparative methods. The phylogenetic comparative approach (Felsenstein, 1985, 2008; Felsenstein and Felenstein, 2004; Grafen, 1989; Harvey and Pagel, 1991; Ridley and Grafen, 1996) allows us to ask questions about how the evolution of a trait occurs through comparison of extant species (although fossil evidence can be incorporated) while taking into account shared ancestry estimated from a phylogeny. We can then ask questions such as what factors (e.g. social or ecological) are associated with the evolution of a trait (e.g. brain size), if that trait evolves directionally, how much common ancestry constrains evolution, and how the evolution of a trait influences speciation rates.

The three major hypotheses of neurocognitive evolution that have been proposed focus on identifying primary factors that have driven differences in brain size and cognitive function across species. The first set of hypotheses suggest that cognition has evolved due to the value of *ecological intelligence*; the ability to find and extract food (Byrne, 1997; Parker and Gibson, 1977), manage high spatiotemporal variation in food resources (Sol et al., 2005), or manage and defend large territories (Clutton-Brock and Harvey, 1980). The second set of hypotheses propose that cognition has evolved primarily due to its value in *social intelligence*; the ability to negotiate and succeed through dominance in large groups (Dunbar, 1998; Whiten and Byrne, 1988) or alternatively the ability to manage positive relationships and social partnerships (Dunbar and Shultz, 2007, 2010; Emery et al., 2007). The third hypothesis, recently proposed to reconcile ecological and social drivers, suggests that cognition evolved to *buffer* individuals against environmental challenges by producing appropriate behavioural responses in new socio-ecological contexts (Allman and Hasenstaub, 1999; Deaner et al., 2003; Sol, 2009).

Each of these hypotheses has been tested using comparative methods and each has found some support. For example, brain size depends on diet in mammals (Eisenberg and Wilson, 1978; Gittleman, 1986; Harvey et al., 1980; MacLean et al., 2014) suggesting a role of ecology. Likewise, brain size and neocortex size are related to social group size (Barton and Dunbar, 1997; Dunbar, 1998; Dunbar and Bever, 1998; Gittleman, 1986; Marino, 1996) and other metrics of social group structure in mammals (reviewed in Dunbar and Shultz, 2007) suggesting that social drivers are also important to the evolution of the brain and cognition. Interestingly, comparison of ecological and social factors in ungulates, showed that relative brain size is influenced by social and ecological factors while relative neocortex size is only influenced by sociality (Shultz and Dunbar, 2006). Finally, species with larger brains have been shown to survive better in novel environments (Sol et al., 2005, 2007, 2008) in support to the cognitive buffer hypothesis (Sol, 2009).

Comparative studies focused on brain size have also been largely criticised (Healy and Rowe, 2007; Lihoreau et al., 2012; Roth et al., 2010a). The high cognitive capacity of small-brained invertebrates, such as bees and ants, suggests that high cognitive capabilities do not require large overall brain size (Chittka and Niven, 2009). Measurements of brain size or brain structure volumes are too coarse grained given that current neuroscience methods enable us to study fine scale brain organisation and function (Healy and Rowe, 2007; Roth et al., 2010a). For instance, cognitive neurosciences have revealed different brain networks and mechanism associated with different cognitive abilities. Thus, instead of studying whole brain or neocortex size, comparative studies should focus on neural circuits and functioning that are known to be involved in the cognitive mechanism of interest when possible (Lihoreau et al., 2012).

Efforts to address the problem that brain size may not be the same as cognitive abilities have been made along two lines of comparative research: (i) spontaneous records of cognition-based behaviours (e.g. innovation) in the wild and (ii) comparative psychology experiments in the lab. The first line of research, also called ‘taxonomical counts of cognition in the wild’ (reviewed in Lefebvre, 2011), enables the study of large samples of “spontaneous” behaviour occurring in the selective environment or at least a natural or semi-natural habitat. This approach has confirmed that relative brain size increases with increased tool use and frequency of innovation in birds (Lefebvre et al., 1997, 2004) and primates (Lefebvre et al., 2004; Reader and Laland, 2002), social learning in primates (Reader and Laland, 2002), or deception in primates (Byrne, 2004).

Taking the second approach, a few studies have begun comparing specific cognitive tasks among a small number of related species that differ in social or ecological conditions. One of the most advanced research programs of this kind, has been conduced on North American corvids (Balda and Kamil, 2002; Balda et al., 1996; Kamil, 1998) using a large number of cognitive tests run in the lab. Corvid species that rely heavily on food storing in the wild, such as Clark’s Nutcrakers (*Nucifraga columbiana*), typically outperform other corvids in tasks requiring spatial cognition (Olson et al., 1995); on the other hand, corvid species that are highly social, such as Pinyon Jays (*Gymnorhinus cyanocephalus*), are better in cognition tasks mimicking social challenges such as those designed to evaluate social learning, behavioural flexibility or transitive inference (Bond et al., 2003, 2007, 2010; Templeton et al., 1999). Studies in primates have similarly addressed how social structure is related to the evolution of cognitive abilities. Comparing species that differ in their degree of sociality, Amici et al. (2008) have shown that species with fission-fusion social organisation outperform species with very stable social groups in cognitive tasks requiring inhibitory control and/or flexibility. Very recently, one of the most accomplished studies merging phylogenetic and experimental cognition methods draws a slightly different picture (MacLean et al., 2014). MacLean and his 57 collaborators realized the feat of gathering cognitive performances of 36 animal species (from birds to rodents to apes) in two problem solving tasks measuring self-control. Their results suggest that the major proximate mechanism underlying the evolution of self control is the absolute brain volume rather than residual brain volume corrected for body mass. They also report a significant relationship between cognitive performance and dietary breadth but not social organization in primates. Thus, this massive comparative cognition study challenges both the proxy of cognition (relative brain size) and the hypothesis (social brain hypothesis) tested in many brain comparative studies and illustrates the danger of over interpreting comparative cognition studies. Continued efforts to link specific cognitive functions to their ecological and social settings present a promising avenue to understand the evolution of cognition while recognizing that different cognitive abilities may evolve under different environmental contexts.

A number of new directions using the comparative method have still not been sufficiently exploited. First and foremost, analyses should begin to compare specific regions of the brain or brain function rather than coarse measures of brain size. The increasing ease of using new technology (e.g. MRI, PET) to measure brain structures, connectivity, and function that are frequently measured in cognitive neurosciences could provide another link between the evolution of cognitive processes and ecological or social factors that influence cognition. Second, only a small range of questions using comparative methods have been addressed (see MacLean et al., 2012 for a review). For example, comparative methods can be used to examine the sequence of events in coevolution such that we could ask if the increase of a cognitive ability generally precedes or succeeds specific social or ecological changes. Likewise, we could examine the relative rates of evolution during the increase or decrease of a particular cognitive ability. Finally, we can ask how shifts in cognition are associated with the speciation process itself (e.g. Nicolakakis et al., 2003). Does the evolution of increased cognitive ability facilitate speciation?

### Intraspecific selection on neurocognitive traits: the fitness approach

Measuring contemporary selection has proved a powerful approach to understanding the evolution of traits and this method could be readily applied to the evolution of cognition. The basic premise of this ‘*fitness*’ approach follows Darwin’s theory of evolution (1859) which suggests that short term selection is the primary cause of evolutionary change and speciation. Therefore a careful examination of selection can help us understand how a trait evolves. Selection can come from a number of origins which largely fall under *natural selection*, which includes the effects of abiotic influences and interspecific interactions on survival and reproduction (Darwin, 1859; and modern synthesis in Huxley, 1942), or *social selection* (Lyon and Montgomerie, 2012; West-Eberhard, 1983), which includes selection due to intraspecific social interactions including the effects of mating behaviour (i.e. sexual selection, Darwin, 1871) and kin cooperation (i.e. kin selection, Hamilton, 1964) among other intraspecific interactions. There are two distinct advantages to the fitness method relative to the comparative method for studying neurocognitive evolution. The first advantage is that studies of selection measure fitness costs and benefits of specific traits which can provide a close match with measurements of cognitive abilities and brain mechanisms currently studied in animal cognition and neurosciences (Figure 2). Thus the fitness approach provides opportunities to integrate our proximate understanding of cognition with new findings on the ultimate causes of cognitive evolution. The second advantage is that examination of selection ideally includes identification of the agent of selection or the specific social or ecological challenges that favour a specific trait. Adopting this approach helps us acknowledge that there may be multiple factors that select for a given cognitive ability and that each species will require only a subset of all cognitive skills given their environment.

**Figure 2:**
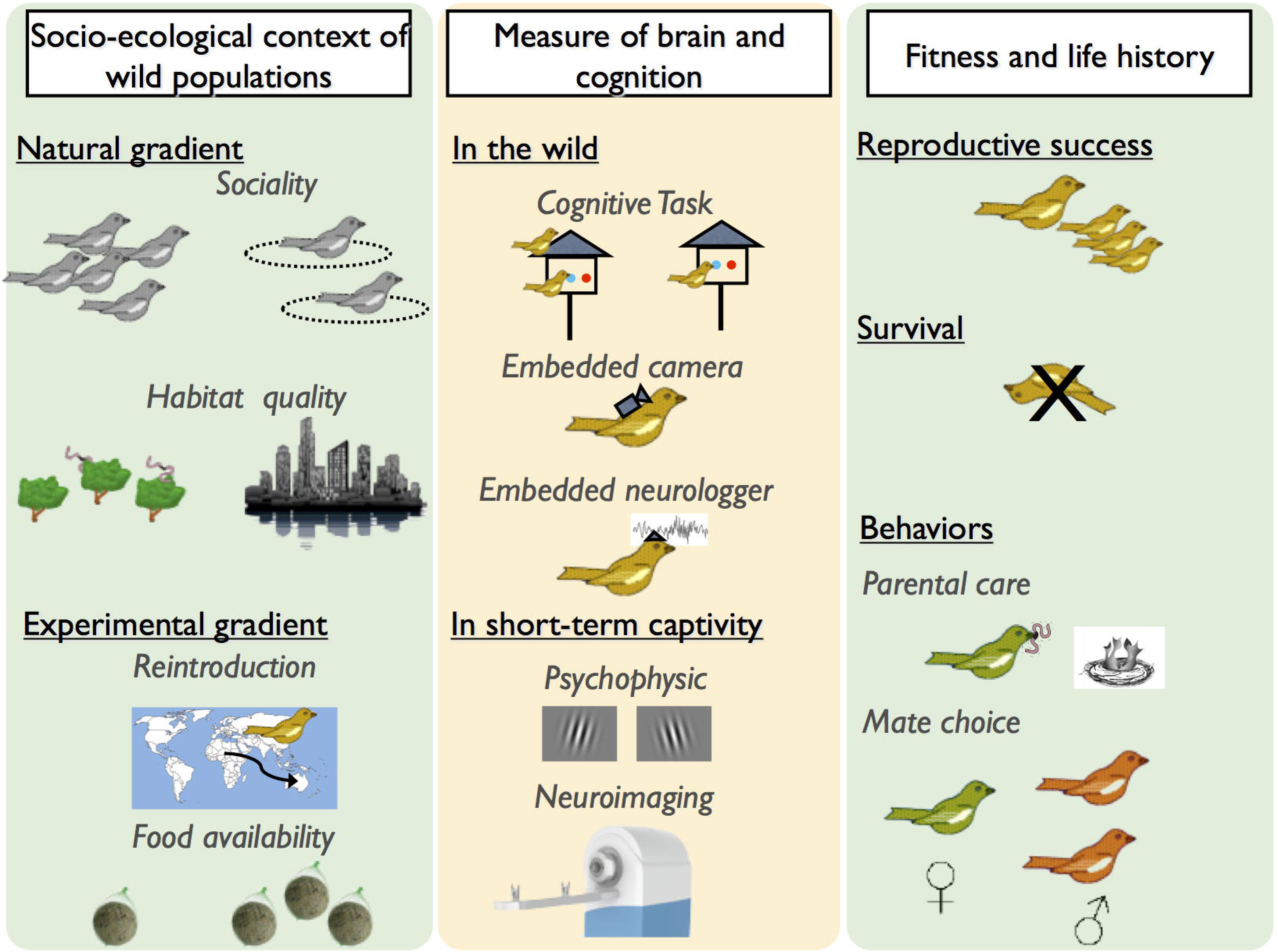
How to study brain and cognition selection? Ideal evolutionary ecology studies of cognition should integrate socio-ecological (left panel, 1), neurocognitive (middle panel, 2) and fitness (right panel, 3) variables. Such an approach seeks to truly merge behavioural and evolutionary (green background) and cognitive neuroscience (yellow background) methods. As examples:

1. Socio-ecological contexts of selection could correspond to natural gradients in sociality (ie: Population density, gregariousness), habitat quality (ie: level of fragmentation, urbanization) and/or distribution of resources (ie: harshness of the environment). Experimental manipulations of ecological factors, such as variation in food supplementation or reintroduction in a novel environment, are of particular interest to isolate ecological causes of selection.
2. Cognitive abilities can be measured in the wild using foraging tasks. This approach has been successfully adapted to measure perception, problem solving, learning, behavioural flexibility and spatial cognition. Such methods rely on individual identification usually mediated by visual tags (i.e. colour rings) or passive integrated transponders (PIT) tags. However, some cognitive functions are difficult to measure in the wild and one may want to have a better control on motivational state and environmental parameters. Short-term period of captivity seems appropriate in such a framework and potentially enable us to use up-to-date psychophysics protocols and equipment developed in comparative cognition labs. Development of embedded cameras or microphones has the potentials to reveal spontaneous cognitive capabilities like tool use, social cognition or vocal communication. Likewise, neurologgers or transmitters enable us to measure brain activity (electroencephalogram, single unit activity) in free ranging wild animals. Spatial and whole brain measurement could also be assessed using MRI or PET devices through short term scanning protocol.
3. The fitness benefit is traditionally assessed through evaluation of reproductive success or a measure of survival. Behaviour associated with reproductive success (i.e. mating, parental care) can also be used as proxies of fitness.

To show that animal cognition evolves under direct natural or social selection requires that the three necessary conditions for selection and evolution that Darwin (1859, 1871) outlined apply to cognitive abilities (Dukas, 2004). Traits, or in this case cognitive abilities, will evolve if (1) there is variability in cognition between individuals, (2) that this variability in cognitive ability is heritable, and (3) that this variation is related to variance in fitness (survival, reproductive success) under specific environmental conditions. Few studies have tackled these questions specifically, but evidence from the literature supports the notion that cognition *should* evolve under selection making the fitness approach fruitful.

#### (1) Variation in neurocognitive ability

Inter-individual variability in animal cognition studies is rarely reported, yet without variation, cognition can not evolve. Studies in animal cognition generally focus on a small number of individuals because of the time involved in training and testing subjects and this small sample size precludes useful estimates of variation in cognitive abilities. However, a recent meta-analysis of variation in individual performances at three common cognitive tasks for different species revealed very high inter-individual variability (Thornton and Lukas, 2012). Individual performances varied almost continuously from 25-100% success at a task in tests for species with the largest sample sizes. Some of this variation is influenced by age, sex, developmental conditions, or previous experience, so determining the extent of variation due to additive genes rather than plasticity will require large sample sizes at single cognitive tasks.

Despite little direct evidence, there are a number of indirect measures of cognitive variability that further support the notion that intraspecific variation in cognitive abilities should be widespread. A growing number of recent studies focus on intraspecific variation in brain size including both within and among population variation (for a review see Gonda et al., 2013). This variation is also apparent in humans where inter-individual variation in brain structure and function has often been considered “noise” until recently (Kanai and Rees, 2011). Perhaps the best evidence of inter-individual cognitive variation comes from research on “general intelligence” in humans, which has been extensively documented through the use of intelligence or ‘IQ’ tests and shows high variation among individuals (Deary et al., 2010). Recent work has sought to tie variation between IQ in humans to its neural substrate (Deary et al., 2010; Penke et al., 2012).

#### (2) Heritability of neurocognitive abilities

Heritability of traits is difficult to measure since many non-genetic effects (common environment, parental care, maternal effects, etc…) contribute to resemblance between parents and offspring. For example, twin studies show that brain structure or function (e.g. face recognition) is heritable in humans (Peper et al., 2007; Wilmer et al., 2010; Zhu et al., 2010), yet non-genetic effects that occur in utero can not be excluded (but see Trzaskowski et al., 2013). One of the most powerful approaches to demonstrate that heritability of cognitive traits exists is through artificial selection experiments where species show phenotypic changes in response to researcher imposed selection criteria. Mery and Kawecki have shown that associative learning abilities for choice of oviposition substrate can be inherited in *Drosophila melanogaster* (see Kawecki, 2010 for a review; Mery and Kawecki, 2002, 2003, 2005). Marked differences in learning and memory were shown between high learning and low learning selected *Drosophila* populations over 15 generations. Artificial selection of brain size in guppies (*Poecilia reticulata*) also suggests heritability of brain size (Kotrschal et al., 2013a) with a divergence in relative brain size of 9% between lines selected for large vs. small size over just two generations. Interestingly, large-brained females outperformed small-brained females in a numerical learning test, which also provides evidence for an association between increased brain size and higher cognition. These results should be treated cautiously since disentangling true heritability from plasticity would require more than 2 generations and a relaxation of selection to see if brain size differences persist (see Healy and Rowe, 2013; Kotrschal et al., 2013b). Finally, the use of genome wide association has recently been used to demonstrate a genetic basis of human general intelligence and cognition. This approach has shown that a substantial proportion (between 40 and 66%) of individual differences in human general intelligence is linked to genetic variation (Davies et al., 2011; Benyamin et al., 2013; but see Chabris et al., 2012; Deary et al., 2012; Plomin et al., 2013).

#### (3) The fitness benefits of cognition

Selection on cognitive abilities will occur if there are fitness benefits to particular cognitive phenotypes under a given set of environmental conditions. Addressing this question is challenging because it requires both an estimate of cognitive performance or brain structure/activity of a large number of individuals as well as fitness estimates, such as reproductive success or survival, for the same individuals. Most cognitive tests are run under laboratory conditions to control confounding effects on cognition and yet the best estimates of fitness benefits should be measured in the wild where the importance of a specific cognitive ability will also depend on the environmental context. Fitness measured in artificial selection experiments on cognition or brain size have reported costs and benefits of improved cognitive abilities in insects (Dukas, 2008; Kawecki, 2010) or increased brain size in fishes (Kotrschal et al., 2013a), but the value of these traits in nature are unknown. In humans, general intelligence is correlated with school achievement, job performance, health, and survival (Deary et al., 2010), but not necessarily actual fitness (i.e. number of lifetime offspring that reproduce).

Two very recent studies have finally succeeded in measuring fitness consequences of problem-solving abilities in wild populations of great tits (*Parus major*) (Cauchard et al., 2012; Cole et al., 2012). Cole et al. (2012) took birds into short term captivity to perform an innovation task to get food. Birds who solved the task had larger clutch sizes, but tended to desert their nest more often if disturbed (Cole et al. 2012). Cauchard et al. (2013) conducted cognitive tests in the wild, where birds had to remove an obstacle that blocked access to their nestbox. Those who could solve the puzzle had higher survival of offspring to fledging. Both studies found individual variation in cognitive performance of birds (solvers vs. non solvers), so selection should act on problem solving abilities. Fitness costs of higher cognition (e.g. higher desertion rates; Cole et al. 2012), could produce a trade-off that helps maintain variation in cognitive abilities among individuals. These results are very promising, and should be diversified to a much broader range of cognitive abilities and expanded to measures of brain structure or function (Figure 2). Furthermore, following pioneering research linking food hoarding behaviour and spatial memory (see Pravosudov and Roth II, 2013 for a review, and see Case Study 1 below), understanding why cognition evolves will also require us to directly link cognitive performance (e.g. memory) to ecological challenges that the animals face in their natural environment (e.g. finding a food store). This last point is critical because if there are correlations among different cognitive abilities then measurement of selection (i.e. higher fitness) on one ability could be due to correlational selection on a different cognitive trait that is the actual target of selection.

### Case Studies

As detailed above, there is now some evidence for selection of cognitive abilities in wild animals, including humans. The next challenge for cognitive ecology is to identify which cognitive functions are critical for a species in their natural environment. While for most species we are still at the point of forming hypotheses on which cognitive abilities are critical (as we did for mate choice in Figure 1), there are a few studies that have moved well beyond this stage. Here we present two lines of research as examples of successfully linking natural behaviour, cognitive function and ecological agents of selection.

#### (1) The evolutionary ecology of spatial memory

Food hoarding animals rely on food caching and later retrieval of caches to survive winter and should have evolved excellent spatial memory abilities and associated neural structures (i.e. hippocampus). Based on this simple ecology-driven hypothesis, a flourishing literature on the cognitive ecology of food storing has emerged over the last thirty years. This work has successfully combined proximate and ultimate understandings of spatial cognition and serves as an example for future studies of the evolutionary ecology of cognition (see Brodin, 2010 for an historical review).

The first studies of the evolutionary and ecological significance of spatial memory employed the comparative framework, with the prediction that scatter food hoarding species should surpass non-hoarding species in spatial memory tasks and should have a relatively bigger hippocampus. However, results from these early studies were equivocal and difficult to interpret. The superiority of spatial capabilities in hoarding species was not always clear (reviewed in Healy et al., 2009). Furthermore, and more concerning, the comparative approach suffers from a number of confounding factors, such as morphological differences between species, that could never clearly be separated from performance in cognitive tasks (but see (Kamil, 1998) for methods).

Problems with comparative analyses have been very elegantly solved by focusing on intra-specific variation in a number of landmark studies comparing populations exposed to different ecological contexts. In one of the earliest of such studies, Pravosudov and Clayton (2002) demonstrated that black-capped chickadees (*Poecile atricapilla*) living in harsh winter climates (i.e. Alaska) cache more food, have higher spatial memory capabilities, and have a larger hippocampus that contains more neurones than individuals of the same species in populations from milder climates (i.e. Colorado). While the appearance of adaptation is clear, such differences could reflect either local adaptation shaped by natural selection or result from plasticity in brain structure and behaviour generated from the local environment. The persistence of among population differences in brain structure and caching behaviour in common garden experiments, during which 10 days-old chicks from these different populations were hand-raised in identical environmental conditions, strongly argues for a role of natural selection in shaping local adaptation for spatial memory, neural density, and neurogenesis in the hippocampus (Roth et al., 2010b, 2012).

Recent analyses using this within species comparative approach in this and other species have further pushed our understanding of the links between cognition and evolutionary ecology and between proximate and ultimate understandings of cognitive evolution. Research in mountain chickadees (*Poecile gambeli*) along an altitudinal gradient has shown similar patterns of differentiation in food storage, spatial memory, and hippocampal characteristics as with contrasted populations in the black-capped chickadee (Freas et al., 2013). Other studies have extended this work on spatial memory differences across populations in caching behaviour to differences between behavioural strategies within a population (LaDage et al., 2013). In side-blotched lizards (*Uta stansburiana*), males adopt one of three different mating strategies that rely to different degrees on spatial memory for territory defence and the distribution of available females across territories. Accordingly, characteristics of the dorsal cortex and hippocampus show differences among genetically determined alternative male strategies within a population (Ladage et al., 2009; LaDage et al., 2013). Work on hippocampal size contrasts among populations has recently been extended by fine scale studies of neural structure (Roth et al., 2010a, 2012) and differential gene expression (Pravosudov et al., 2013) within the hippocampus among contrasted populations of birds. The next step should be to measure the influence of spatial cognition and the underlying hippocampal structures or function on fitness in these contrasted environments.

#### (2) Cognitive mechanisms of host-parasite arm races in brood parasites

Avian brood parasites lay their eggs in the nest of other individuals from the same or different species to avoids the costs of parental care but imposes a cost on the host (reviewed in Davies, 2011). These reciprocal selection pressures have often led to an arms race of detection and mimicry in egg appearance – a true cognitive battleground. Studies of avian brood parasitism provide measures of selection on cognitive traits (recognition, rejection, deception), clear identification of the agent of selection, examination of how cognition influences the coevolutionary arms race, and neural traits associated with host-parasite life history.

Studies of avian brood parasitism have done an outstanding job of quantifying the fitness costs and benefits to each player of the host-parasite arms race—often linked to recognition of parasites (Davies, 2011; Lyon and Eadie, 2008). A parasite’s fitness is so intricately tied to acceptance by hosts that they must adapt to new host defences either by identifying and changing to a new host or surpassing host defences. Hosts, on the other hand, pay a cost of parasitism, but the evolution of new defences (often a cognitive ability) must be balanced against the frequency of parasitism and the costs of producing better defences (Davies and Brooke, 1988, 1989a, 1989b; Lotem, 1993; Lotem et al., 1995; Rothstein, 1982). Costs of new defences include developing the cognitive or morphological structures for new defences as well as the added risk of expressing those defences (e.g. rejecting own eggs), and these costs influence the evolution of recognition abilities. Plasticity in host recognition reveals the importance that making an incorrect choice can have for the evolution of egg rejection. For example, some common cuckoo hosts avoid rejecting their own eggs (recognition error) when parasites are not present by only increasing rejection rates when adult cuckoos are seen in the vicinity of the nest (Davies and Brooke 1988). In South American coots, intraspecific parasitism leads to egg rejection, but an interspecific parasite, the blackheaded duck, that imposes no parental care costs is only rejected when ecological conditions render incubation more costly (Lyon and Eadie, 2004). Globally, studies of avian brood parasites have provided an excellent understanding of the selective environment generated by host-parasite interactions that influences the evolution of recognition and rejection of eggs.

Mimicry-recognition-rejection arms races reveal the link between cognitive abilities and the evolutionary dynamics of host-parasite systems. Arms races in avian brood parasites related to egg mimicry push host recognition systems to identify parasites while avoiding recognition errors (Davies and Brooke, 1988; Rothstein, 1982). The accuracy of identifying a mimetic egg depends on visual discrimination abilities and recent studies have begun to specifically integrate this process using ‘visual modelling’–information on cone sensitivity and objective measures of egg colour patterns–to understand rejection behaviour, or the lack thereof, in some species (Cassey et al., 2008; Spottiswoode and Stevens, 2010). Recent findings show that visual detection of parasites can improve by integrating multiple sources of information (Spottiswoode and Stevens 2010). Egg cues (de la Colina et al., 2012; Langmore et al., 2009; Spottiswoode and Stevens, 2010; Svennungsen and Holen, 2010), external cues of parasite presence (Davies and Brooke, 1988), or counting the number of eggs laid (Lyon, 2003) have all been shown as means to improve the decision to reject parasite eggs. Use of multiple and disparate cues to improve the accuracy of rejection behaviour would require executive functions to weigh these different criteria in a rejection decision and future research could examine this cognitive ability. Not all host species reject eggs or chicks, which implies that physiological or cognitive limitations may also influence the detection of a parasitic egg (Davies and Brooke, 1988; Lotem, 1993; Lotem et al., 1995; Rodríguez-Gironés and Lotem, 1999; Rothstein, 1982).

An understanding of the cognitive mechanisms underlying rejection have also played an important role in understanding why despite close visual mimicry in eggs, nestlings are rarely mimetic. One hypothesis is that unlike egg recognition where comparisons between multiple host eggs and a single parasitic egg makes discrimination possible, having only a single parasite chick in the nest (e.g. common cuckoos) could have severe long term fitness costs if birds *learn* the appearance of their chicks (Lotem, 1993). Indeed, learning does seem to play a role in identification and discrimination of eggs (Rothstein, 1974, 1978; Strausberger and Rothstein, 2009) and possibly chicks (Colombelli-Négrel et al., 2012; Shizuka and Lyon, 2010). A possible solution in some species, such as the North American coot, might be to use extra cues such as hatch order and soft rejection (e.g. lower feeding) to help identify parasitic chicks while reducing the risk of mis-imprinting (Shizuka and Lyon, 2010, 2011). These models and empirical results show that the cognitive mechanisms underlying how a species is able to recognize its eggs and chicks plays an important role in the evolution of the host-parasite arms race.

Finally, a few studies have also begun to investigate the link between neurophysiology and the ecology of brood parasites. Initial studies focused primarily on whole brain size or hippocampus size in brood parasites and their non-parasitic relatives since each species should face different ecological imperatives. Generally, whole brain size tends to be smaller in brood-parasites than their closest relatives (Corfield et al., 2013; Iwaniuk, 2004; Overington, 2011), which could be linked to less complex cognitive function needed in the absence of parental care in brood parasites (Boerner and Krüger, 2008). Hippocampus size, however, varies predictably with the need for excellent spatial memory in brood parasites. Brood parasites have an enlarged hippocampus in the breeding season (Clayton et al., 1997), the sex that searches for nests tends to have a larger hippocampus than the other sex (Reboreda et al., 1996; Sherry et al., 1993), and brood parasites have a relatively larger hippocampus than closely related non-parasites (Corfield et al., 2013; Reboreda et al., 1996). Furthermore, recent analysis has uncovered a specific region of the hippocampus that is enlarged in parasitic species relative to others (Nair-Roberts et al., 2006), suggesting brain regions may have evolved to manage the specific needs of brood parasites relative to other spatial memory. These studies provide a rare example of direct linkage between ecology and neurophysiology on a well understood fitness landscape. An exciting next step in such systems could be to examine variation in neural structure with variation in the ability of different hosts – either across or within a species – to reject parasitic eggs or chicks.

The above studies provide some of the best examples of how discrimination ability links with cognitive decision making under natural ecological conditions. While many of these host-parasite studies have not specifically been framed in terms of cognitive ecology, the focus on discrimination, recognition, learning, and decision making are all clearly linked to cognition and could further link to both specific cognitive abilities studied in other organisms and to neurophysiological studies. Together with the strong understanding of the fitness costs and benefits of host-parasite coevolution, these systems provide an excellent opportunity to link cognition, neurophysiology, and evolutionary biology.

## Conclusion

We have highlighted two ways to investigate the evolution of cognitive processes in animals: the comparative approach focuses on evolutionary history while the fitness approach examines contemporary selection. Much of our knowledge on the evolution of cognition comes from the comparative approach and the full application of recently developed phylogenetic tools should allow for interesting new results in this line of research. However, since cognition presents all the characteristics of traits under selection (variation, heritability and fitness benefits), we believe that taking the fitness approach to cognitive function will allow us to better explore the evolutionary mechanisms that shape animal minds. Furthermore, the fitness approach more easily allows us to integrate proximate and ultimate factors underlying animal cognition in a single study, as suggested fifty years ago by Tinbergen (Tinbergen, 1963).

## 4- Future directions

The integration of evolutionary biology with cognitive sciences provides a very promising avenue of research that could revolutionize our understanding of animal mind. Here we highlighted how methods and new research questions in evolutionary biology could contribute to our current understanding of the proximate basis of cognition. We believe the (unranked) top priorities for the future are:

1) Identify cognitive functions that are crucial for species currently studied by evolutionary ecologists and behavioural ecologists.
2) What are the fitness consequences of cognitive performance in the wild and what are the ecological contexts under which that ability is favoured?
3) Are cognitive performance and/or neurocognitive processes consistent across different environments for a given species? Are they consistent for a given individual if we can measure cognitive abilities in the wild?
4) Can we create more ecologically relevant cognitive performance tasks that help link cognitive abilities or brain structure to specific ecological challenges?
5) What environmental or social factors are associated with the evolution of specific cognitive abilities or neural structures across species and what role do these abilities play in the speciation process?
6) Are different cognitive abilities related to each other (i.e. positive correlation or trade-off)? Is there compelling evidence for general intelligence in non-human animals?
7) Problem solving is the one “cognitive” task that has been related to fitness in wild animals. However the cognitive mechanisms underlying this task remain unclear (Healy, 2012; Rowe and Healy, 2014; Thornton et al., 2014). What are the fitness benefits of other well characterized cognitive capacities such as visual cognition or associative learning?
8) What are the implications of cognitive performance for theory in evolutionary ecology and conversely what does an ecological perspective on cognition tell us about neurocognitive development?

## Acknowledgments

This work has been supported by a Fyssen postdoctoral fellowship.

## Bibliography

Allman, J., and Hasenstaub, A. (1999). Brains, maturation times, and parenting★. Neurobiol. Aging 20, 447–454.

Amici, F., Aureli, F., and Call, J. (2008). Fission-fusion dynamics, behavioral flexibility, and inhibitory control in primates. Curr. Biol. 18, 1415–1419.

Balda, R.P., and Kamil, A.C. (2002). Spatial and social cognition in corvids: an evolutionary approach. Cogn. Anim. Empir. Theor. Perspect. Anim. Cogn. 129–134.

Balda, R.P., Kamil, A.C., and Bednekoff, P.A. (1996). Predicting cognitive capacity from natural history. In Current Ornithology, (Springer), pp. 33–66.

Barsalou, L.W. (2014). Cognitive psychology: An overview for cognitive scientists (Psychology Press).

Barton, R.A., and Dunbar, R.I. (1997). 9 Evolution of the social brain. Machiavellian Intell. II Ext. Eval. 2, 240.

Bateson, P., and Laland, K.N. (2013). Tinbergen's four questions: an appreciation and an update. Trends Ecol. Evol. 28, 712–718.

Benyamin, B., Pourcain, Bs., Davis, O.S., Davies, G., Hansell, N.K., Brion, M.-J., Kirkpatrick, R.M., RAM Cents, S.F., Miller, M.B., and Haworth, C.M.A. (2013). Childhood intelligence is heritable, highly polygenic and associated with FNBP1L. Mol. Psychiatry.

Boerner, M., and Krüger, O. (2008). Why do parasitic cuckoos have small brains? Insights from evolutionary sequence analyses. Evol. Int. J. Org. Evol. 62, 3157–3169.

Bond, A.B., and Kamil, A.C. (2002). Visual predators select for crypticity and polymorphism in virtual prey. Nature 415, 609–613.

Bond, A.B., and Kamil, A.C. (2006). Spatial heterogeneity, predator cognition, and the evolution of color polymorphism in virtual prey. Proc. Natl. Acad. Sci. U. S. A. 103, 3214–3219.

Bond, A.B., Kamil, A.C., and Balda, R.P. (2003). Social complexity and transitive inference in corvids. Anim. Behav. 65, 479–487.

Bond, A.B., Kamil, A.C., and Balda, R.P. (2007). Serial reversal learning and the evolution of behavioral flexibility in three species of North American corvids (Gymnorhinus cyanocephalus, Nucifraga columbiana, Aphelocoma californica). J. Comp. Psychol. 121, 372.

Bond, A.B., Wei, C.A., and Kamil, A.C. (2010). Cognitive representation in transitive inference: a comparison of four corvid species. Behav. Processes 85, 283–292.

Boogert, N.J., Fawcett, T.W., and Lefebvre, L. (2011). Mate choice for cognitive traits: a review of the evidence in nonhuman vertebrates. Behav. Ecol. 22, 447–459.

Brodin, A. (2010). The history of scatter hoarding studies. Philos. Trans. R. Soc. B Biol. Sci. 365, 869–881.

Byrne, R.W. (1997). ii The Technical Intelligence hypothesis: An additional evolutionary stimulus to intelligence? Machiavellian Intell. II Ext. Eval. 2, 289.

Byrne, R.W. (2004). Neocortex size predicts deception rate in primates. Proc. R. Soc. B Biol. Sci. 271, 1693.

Cassey, P., Honza, M., Grim, T., and Hauber, M.E. (2008). The modelling of avian visual perception predicts behavioural rejection responses to foreign egg colours. Biol. Lett. 4, 515–517.

Cauchard, L., Boogert, N.J., Lefebvre, L., Dubois, F., and Doligez, B. (2012). Problem-solving performance is correlated with reproductive success in a wild bird population. Anim. Behav.

Chabris, C.F., Hebert, B.M., Benjamin, D.J., Beauchamp, J., Cesarini, D., van der Loos, M., Johannesson, M., Magnusson, P.K., Lichtenstein, P., and Atwood, C.S. (2012). Most reported genetic associations with general intelligence are probably false positives. Psychol. Sci. 23, 1314–1323.

Chittka, L., and Niven, J. (2009). Are bigger brains better? Curr. Biol. 19, R995–R1008.

Clayton, N.S., Reboreda, J.C., and Kacelnik, A. (1997). Seasonal changes of hippocampus volume in parasitic cowbirds. Behav. Processes 41, 237–243.

Clutton-Brock, T.H., and Harvey, P.H. (1980). Primates, brains and ecology. J. Zool. 190, 309– 323.

Cole, E.F., Morand-Ferron, J., Hinks, A.E., and Quinn, J.L. (2012). Cognitive ability influences reproductive life history variation in the wild. Curr. Biol.

De la Colina, M.A., Pompilio, L., Hauber, M.E., Reboreda, J.C., and Mahler, B. (2012). Different recognition cues reveal the decision rules used for egg rejection by hosts of a variably mimetic avian brood parasite. Anim. Cogn. 15, 881–889.

Colombelli-Négrel, D., Hauber, M.E., Robertson, J., Sulloway, F.J., Hoi, H., Griggio, M., and Kleindorfer, S. (2012). Embryonic learning of vocal passwords in superb fairy-wrens reveals intruder cuckoo nestlings. Curr. Biol. CB 22, 2155–2160.

Corfield, J.R., Birkhead, T.R., Spottiswoode, C.N., Iwaniuk, A.N., Boogert, N.J., Gutiérrez-Ibáñez, C., Overington, S.E., Wylie, D.R., and Lefebvre, L. (2013). Brain Size and Morphology of the Brood-Parasitic and Cerophagous Honeyguides (Aves: Piciformes). Brain. Behav. Evol. 81, 170–186.

Darwin, C. (1859). On the origin of the species by natural selection.

Darwin, C. (1871). Sexual selection and the descent of man. Murray Lond.

Davies, N.B. (2011). Cuckoos, cowbirds and other cheats (A&C Black).

Davies, N.B., and Brooke, M. de L. (1988). Cuckoos versus reed warblers: adaptations and counteradaptations. Anim. Behav. 36, 262–284.

Davies, N.B., and Brooke, M. de L. (1989a). An experimental study of co-evolution between the cuckoo, Cuculus canorus, and its hosts. I. Host egg discrimination. J. Anim. Ecol. 207–224.

Davies, N.B., and Brooke, M. de L. (1989b). An experimental study of co-evolution between the cuckoo, Cuculus canorus, and its hosts. II. Host egg markings, chick discrimination and general discussion. J. Anim. Ecol. 225–236.

Davies, G., Tenesa, A., Payton, A., Yang, J., Harris, S.E., Liewald, D., Ke, X., Le Hellard, S., Christoforou, A., and Luciano, M. (2011). Genome-wide association studies establish that human intelligence is highly heritable and polygenic. Mol. Psychiatry 16, 996–1005.

Deaner, R.O., Barton, R.A., and van Schaik, C.P. (2003). 10 Primate Brains and Life Histories: Renewing the Connection. Primate Life Hist. Socioecology 233.

Deary, I.J., Penke, L., and Johnson, W. (2010). The neuroscience of human intelligence differences. Nat. Rev. Neurosci. 11, 201–211.

Deary, I.J., Yang, J., Davies, G., Harris, S.E., Tenesa, A., Liewald, D., Luciano, M., Lopez, L.M., Gow, A.J., and Corley, J. (2012). Genetic contributions to stability and change in intelligence from childhood to old age. Nature 482, 212–215.

Dukas, R. (2004). Evolutionary biology of animal cognition. Annu. Rev. Ecol. Evol. Syst. 347–374.

Dukas, R. (2008). Evolutionary biology of insect learning. Annu Rev Entomol 53, 145–160.

Dukas, R., and Ratcliffe, J.M. (2009). Cognitive ecology II (University of Chicago Press).

Dunbar, R.I. (1998). The social brain hypothesis. Brain 9, 10.

Dunbar, R.I., and Shultz, S. (2007). Evolution in the social brain. Science 317, 1344–1347.

Dunbar, R.I., and Shultz, S. (2010). Bondedness and sociality. Behaviour 147, 775–803.

Dunbar, R.I.M., and Bever, J. (1998). Neocortex size predicts group size in carnivores and some insectivores. Ethology 104, 695–708.

Eisenberg, J.F., and Wilson, D.E. (1978). Relative brain size and feeding strategies in the Chiroptera. Evolution 740–751.

Emery, N.J., Seed, A.M., von Bayern, A.M., and Clayton, N.S. (2007). Cognitive adaptations of social bonding in birds. Philos. Trans. R. Soc. B Biol. Sci. 362, 489–505.

Felsenstein, J. (1985). Phylogenies and the comparative method. Am. Nat. 1–15.

Felsenstein, J. (2008). Comparative methods with sampling error and within-species variation: contrasts revisited and revised. Am. Nat. 171, 713–725.

Felsenstein, J., and Felenstein, J. (2004). Inferring phylogenies (Sinauer Associates Sunderland).

Freas, C.A., Roth, T.C., LaDage, L.D., and Pravosudov, V.V. (2013). Hippocampal neuron soma size is associated with population differences in winter climate severity in food-caching chickadees. Funct. Ecol. 27, 1341–1349.

Giraldeau, L.-A. (2004). Introduction: Ecology and the Central Nervous System. Brain. Behav. Evol. 63, 193–196.

Gittleman, J.L. (1986). Carnivore brain size, behavioral ecology, and phylogeny. J. Mammal. 23–36.

Gonda, A., Herczeg, G., and Merilä, J. (2013). Evolutionary ecology of intraspecific brain size variation: a review. Ecol. Evol. 3, 2751–2764.

Grafen, A. (1989). The phylogenetic regression. Philos. Trans. R. Soc. Lond. B. Biol. Sci. 119–157.

Hamilton, W.D. (1964). The genetical evolution of social behaviour. II. J. Theor. Biol. 7, 17–52.

Harvey, P.H., and Pagel, M. (1991). Comparative method in evolutionary biology (POD).

Harvey, P.H., Clutton-Brock, T.H., and Mace, G.M. (1980). Brain size and ecology in small mammals and primates. Proc. Natl. Acad. Sci. 77, 4387–4389.

Healy, S.D. (2012). Animal Cognition: The Trade-off to Being Smart. Curr. Biol. 22, R840– R841.

Healy, S., and Braithwaite, V. (2000). Cognitive ecology: a field of substance? Trends Ecol. Evol. 15, 22–26.

Healy, S.D., and Rowe, C. (2007). A critique of comparative studies of brain size. Proc. R. Soc. B Biol. Sci. 274, 453–464.

Healy, S.D., and Rowe, C. (2013). Costs and benefits of evolving a larger brain: doubts over the evidence that large brains lead to better cognition. Anim. Behav.

Healy, S.D., Bacon, I.E., Haggis, O., Harris, A.P., and Kelley, L.A. (2009). Explanations for variation in cognitive ability: behavioural ecology meets comparative cognition. Behav. Processes 80, 288–294.

Heyes, C.M., and Huber, L. (2000). The evolution of cognition [electronic resource] (The MIT Press).

Huxley, J. (1942). Evolution. The Modern Synthesis. Evol. Mod. Synth.

Iwaniuk, A.N. (2004). Brood Parasitism and Brain Size in Cuckoos: A Cautionary Tale on the Use of Modern Comparative Methods. Int. J. Comp. Psychol. 17.

Kamil, A.C. (1998). On the proper definition of cognitive ethology. Anim. Cogn. Nat. Acad. Press San Diego 1–28.

Kanai, R., and Rees, G. (2011). The structural basis of inter-individual differences in human behaviour and cognition. Nat. Rev. Neurosci. 12, 231–242.

Kawecki, T.J. (2010). Evolutionary ecology of learning: insights from fruit flies. Popul. Ecol. 52, 15–25.

Kotrschal, A., Rogell, B., Bundsen, A., Svensson, B., Zajitschek, S., Brännström, I., Immler, S., Maklakov, A.A., and Kolm, N. (2013a). Artificial selection on relative brain size in the guppy reveals costs and benefits of evolving a larger brain. Curr. Biol.

Kotrschal, A., Rogell, B., Bundsen, A., Svensson, B., Zajitschek, S., Brännström, I., Immler, S., Maklakov, A.A., and Kolm, N. (2013b). The benefit of evolving a larger brain: big-brained guppies perform better in a cognitive task. Anim. Behav. 86, e4.

Ladage, L.D., Riggs, B.J., Sinervo, B., and Pravosudov, V.V. (2009). Dorsal cortex volume in male side-blotched lizards (Uta stansburiana) is associated with different space use strategies. Anim. Behav. 78, 91–96.

LaDage, L.D., Maged, R.M., Forney, M.V., Roth, T.C., 2nd, Sinervo, B., and Pravosudov, V.V. (2013). Interaction between territoriality, spatial environment, and hippocampal neurogenesis in male side-blotched lizards. Behav. Neurosci. 127, 555–565.

Laland, K.N., Sterelny, K., Odling-Smee, J., Hoppitt, W., and Uller, T. (2011). Cause and effect in biology revisited: is Mayr's proximate-ultimate dichotomy still useful? Science 334, 1512– 1516.

Langmore, N.E., Cockburn, A., Russell, A.F., and Kilner, R.M. (2009). Flexible cuckoo chick-rejection rules in the superb fairy-wren. Behav. Ecol. arp086.

Lefebvre, L. (2011). Taxonomic counts of cognition in the wild. Biol. Lett. 7, 631–633.

Lefebvre, L., Whittle, P., Lascaris, E., and Finkelstein, A. (1997). Feeding innovations and forebrain size in birds. Anim. Behav. 53, 549–560.

Lefebvre, L., Reader, S.M., and Sol, D. (2004). Brains, innovations and evolution in birds and primates. Brain. Behav. Evol. 63, 233–246.

Lihoreau, M., Latty, T., and Chittka, L. (2012). An exploration of the social brain hypothesis in insects. Front. Physiol. 3.

Lotem, A. (1993). Learning to recognize nestlings is maladaptive for cuckoo Cuculus canorus hosts. Nature 362, 743–745.

Lotem, A., Nakamura, H., and Zahavi, A. (1995). Constraints on egg discrimination and cuckoo-host co-evolution. Anim. Behav. 49, 1185–1209.

Lyon, B.E. (2003). Egg recognition and counting reduce costs of avian conspecific brood parasitism. Nature 422, 495–499.

Lyon, B.E., and Eadie, J.M. (2004). An obligate brood parasite trapped in the intraspecific arms race of its hosts. Nature 432, 390–393.

Lyon, B.E., and Eadie, J.M. (2008). Conspecific brood parasitism in birds: a life-history perspective. Annu. Rev. Ecol. Evol. Syst. 39, 343–363.

Lyon, B.E., and Montgomerie, R. (2012). Sexual selection is a form of social selection. Philos. Trans. R. Soc. B Biol. Sci. 367, 2266–2273.

MacLean, E.L., Matthews, L.J., Hare, B.A., Nunn, C.L., Anderson, R.C., Aureli, F., Brannon, E.M., Call, J., Drea, C.M., and Emery, N.J. (2012). How does cognition evolve? Phylogenetic comparative psychology. Anim. Cogn. 15, 223–238.

MacLean, E.L., Hare, B., Nunn, C.L., Addessi, E., Amici, F., Anderson, R.C., Aureli, F., Baker, J.M., Bania, A.E., and Barnard, A.M. (2014). The evolution of self-control. Proc. Natl. Acad. Sci. 201323533.

Marino, L. (1996). What can dolphins tell us about primate evolution? Evol. Anthropol. Issues News Rev. 5, 81–86.

Mery, F., and Kawecki, T.J. (2002). Experimental evolution of learning ability in fruit flies. Proc. Natl. Acad. Sci. 99, 14274–14279.

Mery, F., and Kawecki, T.J. (2003). A fitness cost of learning ability in Drosophila melanogaster. Proc. R. Soc. Lond. B Biol. Sci. 270, 2465–2469.

Mery, F., and Kawecki, T.J. (2005). A cost of long-term memory in Drosophila. Science 308, 1148–1148.

Nair-Roberts, R.G., Erichsen, J.T., Reboreda, J.C., and Kacelnik, A. (2006). Distribution of substance P reveals a novel subdivision in the hippocampus of parasitic South American cowbirds. J. Comp. Neurol. 496, 610–626.

Nicolakakis, N., Sol, D., and Lefebvre, L. (2003). Behavioural flexibility predicts species richness in birds, but not extinction risk. Anim. Behav. 65, 445–452.

Olson, D.J., Kamil, A.C., Balda, R.P., and Nims, P.J. (1995). Performance of four-seed caching corvid species in operant tests of nonspatial and spatial memory. J. Comp. Psychol. 109, 173.

Overington, S.E. (2011). Behavioural innovation and the evolution of cognition in birds.

Parker, S.T., and Gibson, K.R. (1977). Object manipulation, tool use and sensorimotor intelligence as feeding adaptations in Cebus monkeys and great apes. J. Hum. Evol. 6, 623–641.

Penke, L., Maniega, S.M., Bastin, M.E., Hernández, M.V., Murray, C., Royle, N.A., Starr, J.M., Wardlaw, J.M., and Deary, I.J. (2012). Brain white matter tract integrity as a neural foundation for general intelligence. Mol. Psychiatry 17, 1026–1030.

Peper, J.S., Brouwer, R.M., Boomsma, D.I., Kahn, R.S., Pol, H., and Hilleke, E. (2007). Genetic influences on human brain structure: a review of brain imaging studies in twins. Hum. Brain Mapp. 28, 464–473.

Plomin, R., Haworth, C.M., Meaburn, E.L., Price, T.S., and Davis, O.S. (2013). Common DNA markers can account for more than half of the genetic influence on cognitive abilities. Psychol. Sci. 24, 562–568.

Pravosudov, V., and Roth II, T.C. (2013). Food Hoarding and the Evolution of Spatial Memory and the Hippocampus. Annu. Rev. Ecol. Evol. Syst. 44.

Pravosudov, V.V., and Clayton, N.S. (2002). A test of the adaptive specialization hypothesis: population differences in caching, memory, and the hippocampus in black-capped chickadees (Poecile atricapilla). Behav. Neurosci. 116, 515.

Pravosudov, V.V., and Smulders, T.V. (2010). Integrating ecology, psychology and neurobiology within a food-hoarding paradigm. Philos. Trans. R. Soc. B Biol. Sci. 365, 859–867.

Pravosudov, V.V., Roth, T.C., 2nd, Forister, M.L., Ladage, L.D., Kramer, R., Schilkey, F., and van der Linden, A.M. (2013). Differential hippocampal gene expression is associated with climate-related natural variation in memory and the hippocampus in food-caching chickadees. Mol. Ecol. 22, 397–408.

Reader, S.M., and Laland, K.N. (2002). Social intelligence, innovation, and enhanced brain size in primates. Proc. Natl. Acad. Sci. 99, 4436–4441.

Real, L.A. (1993). Toward a cognitive ecology. Trends Ecol. Evol. 8, 413–417.

Reboreda, J.C., Clayton, N.S., and Kacelnik, A. (1996). Species and sex differences in hippocampus size in parasitic and non-parasitic cowbirds. Neuroreport 7, 505–508.

Ridley, M., and Grafen, A. (1996). How to study discrete comparative methods. Phylogenies Comp. Method Anim. Behav. 70–103.

Rodríguez-Gironés, M.A., and Lotem, A. (1999). How to detect a cuckoo egg: a signal-detection theory model for recognition and learning. Am. Nat. 153, 633–648.

Roth, T.C., Brodin, A., Smulders, T.V., LaDage, L.D., and Pravosudov, V.V. (2010a). Is bigger always better? A critical appraisal of the use of volumetric analysis in the study of the hippocampus. Philos. Trans. R. Soc. B Biol. Sci. 365, 915–931.

Roth, T.C., LaDage, L.D., and Pravosudov, V.V. (2010b). Learning capabilities enhanced in harsh environments: a common garden approach. Proc. R. Soc. B Biol. Sci. 277, 3187–3193.

Roth, T.C., LaDage, L.D., Freas, C.A., and Pravosudov, V.V. (2012). Variation in memory and the hippocampus across populations from different climates: a common garden approach. Proc. R. Soc. B Biol. Sci. 279, 402–410.

Rothstein, S.I. (1974). Mechanisms of avian egg recognition: possible learned and innate factors. The Auk 796–807.

Rothstein, S.I. (1978). Mechanisms of avian egg-recognition: additional evidence for learned components. Anim. Behav. 26, 671–677.

Rothstein, S.I. (1982). Mechanisms of avian egg recognition: Which egg parameters elicit responses by rejecter species? Behav. Ecol. Sociobiol. 11, 229–239.

Rowe (1999). Receiver psychology and the evolution of multicomponent signals. Anim. Behav. 58, 921–931.

Rowe, C. (2013). Receiver psychology: a receiver's perspective. Anim. Behav. 85, 517–523.

Rowe, C., and Healy, S.D. (2014). Measuring variation in cognition. Behav. Ecol. aru090.

Sherry, D.F., Forbes, M.R., Khurgel, M., and Ivy, G.O. (1993). Females have a larger hippocampus than males in the brood-parasitic brown-headed cowbird. Proc. Natl. Acad. Sci. 90, 7839–7843.

Shettleworth, S.J. (2010). Cognition, evolution, and behavior 2nd edition New York: Oxford University Press.

Shizuka, D., and Lyon, B.E. (2010). Coots use hatch order to learn to recognize and reject conspecific brood parasitic chicks. Nature 463, 223–226.

Shizuka, D., and Lyon, B.E. (2011). Hosts Improve the Reliability of Chick Recognition by Delaying the Hatching of Brood Parasitic Eggs. Curr. Biol. 21, 515–519.

Shultz, S., and Dunbar, R.I.M. (2006). Both social and ecological factors predict ungulate brain size. Proc. R. Soc. B Biol. Sci. 273, 207–215.

Sol, D. (2009). Revisiting the cognitive buffer hypothesis for the evolution of large brains. Biol. Lett. 5, 130–133.

Sol, D., Duncan, R.P., Blackburn, T.M., Cassey, P., and Lefebvre, L. (2005). Big brains, enhanced cognition, and response of birds to novel environments. Proc. Natl. Acad. Sci. U. S. A. 102, 5460–5465.

Sol, D., Székely, T., Liker, A., and Lefebvre, L. (2007). Big-brained birds survive better in nature. Proc. R. Soc. B Biol. Sci. 274, 763–769.

Sol, D., Bacher, S., Reader, S.M., and Lefebvre, L. (2008). Brain size predicts the success of mammal species introduced into novel environments. Am. Nat. 172, S63–S71.

Spottiswoode, C.N., and Stevens, M. (2010). Visual modeling shows that avian host parents use multiple visual cues in rejecting parasitic eggs. Proc. Natl. Acad. Sci. 107, 8672–8676.

Strausberger, B.M., and Rothstein, S.I. (2009). Parasitic cowbirds may defeat host defense by causing rejecters to misimprint on cowbird eggs. Behav. Ecol. 20, 691–699.

Svennungsen, T.O., and Holen, Ø.H. (2010). Avian brood parasitism: information use and variation in egg-rejection behavior. Evol. Int. J. Org. Evol. 64, 1459–1469.

Templeton, J.J., Kamil, A.C., and Balda, R.P. (1999). Sociality and social learning in two species of corvids: the pinyon jay (Gymnorhinus cyanocephalus) and the Clark's nutcracker (Nucifraga columbiana). J. Comp. Psychol. 113, 450.

Théry, M., and Casas, J. (2002). Predator and prey views of spider camouflage. Nature 415, 133.

Thornton, A., and Lukas, D. (2012). Individual variation in cognitive performance: developmental and evolutionary perspectives. Philos. Trans. R. Soc. B Biol. Sci. 367, 2773–2783.

Thornton, A., Clayton, N.S., and Grodzinski, U. (2012). Animal minds: from computation to evolution. Philos. Trans. R. Soc. B Biol. Sci. 367, 2670–2676.

Thornton, A., Isden, J., and Madden, J.R. (2014). Toward wild psychometrics: linking individual cognitive differences to fitness. Behav. Ecol. aru095.

Tinbergen, N. (1963). On aims and methods of ethology. Z. Für Tierpsychol. 20, 410–433.

Trzaskowski, M., Davis, O.S., DeFries, J.C., Yang, J., Visscher, P.M., and Plomin, R. (2013). DNA Evidence for Strong Genome-Wide Pleiotropy of Cognitive and Learning Abilities. Behav. Genet. 1–7.

West-Eberhard, M.J. (1983). Sexual selection, social competition, and speciation. Q. Rev. Biol. 155–183.

Whiten, A., and Byrne, R.W. (1988). Tactical deception in primates. Behav. Brain Sci. 11, 233– 244.

Wilmer, J.B., Germine, L., Chabris, C.F., Chatterjee, G., Williams, M., Loken, E., Nakayama, K., and Duchaine, B. (2010). Human face recognition ability is specific and highly heritable. Proc. Natl. Acad. Sci. 107, 5238–5241.

Zhu, Q., Song, Y., Hu, S., Li, X., Tian, M., Zhen, Z., Dong, Q., Kanwisher, N., and Liu, J. (2010). Heritability of the specific cognitive ability of face perception. Curr. Biol. 20, 137–142.

